# A comparative assessment of adult mosquito trapping methods to estimate spatial patterns of abundance and community composition in southern Africa

**DOI:** 10.1101/633552

**Authors:** Erin E. Gorsich, Brianna R. Beechler, Peter M. van Bodegom, Danny Govender, Milehna M. Guarido, Marietjie Venter, Maarten Schrama

**Affiliations:** Institute of Environmental Sciences, Leiden University, Leiden, NL; School of Life Sciences, University of Warwick, Coventry, UK; The Zeeman Institute for Systems Biology & Infectious Disease Epidemiology Research, University of Warwick, Coventry, UK; College of Veterinary Medicine, Oregon State University, Corvallis, OR, USA; SANPARKS, Scientific Services, Skukuza, South Africa; Zoonotic Arbo- and Respiratory Virus Program, Centre for Viral Zoonoses, Faculty of Health Sciences, University of Pretoria, Pretoria, South Africa

**Keywords:** Arboviruses, Community composition, Kruger National Park, Mosquito, South Africa, Trap bias, Vector

## Abstract

**Background:** Assessing adult mosquito populations is an important component of disease surveillance programs and ecosystem health assessments. Inference from adult trapping datasets involves comparing populations across space and time, but comparisons based on different trapping methods may be biased if traps have different efficiencies or sample different subsets of the mosquito community.

**Methods:** We compared four widely-used trapping methods for adult mosquito data collection in Kruger National Park (KNP), South Africa: Centers for Disease Control miniature light trap (CDC), Biogents Sentinel trap (BG), Biogents gravid *Aedes* trap (GAT) and a net trap. We quantified how trap choice and sampling effort influence inferences on the regional distribution of mosquito abundance, richness and community composition.

**Results:** The CDC and net traps together collected 96% (47% and 49% individually) of the 955 female mosquitoes sampled and 100% (85% and 78% individually) of the 40 species or species complexes identified. The CDC and net trap also identified similar regional patterns of community composition. However, inference on the regional patterns of abundance differed between these traps because mosquito abundance in the net trap was influenced by variation in weather conditions. The BG and GAT traps collected significantly fewer mosquitoes, limiting regional comparisons of abundance and community composition.

**Conclusions:** This study represents the first systematic assessment of trapping methods in natural savanna ecosystems in southern Africa. We recommend the CDC trap or the net trap for future monitoring and surveillance programs.

## Background

Adult mosquito sampling is a key component of mosquito surveillance [1–4], but trapping success may vary across studies due to differences in trapping methods. Different traps vary in their ability to catch certain species and life stages [5–8]. For example, the dominant species attracted with light-baited traps may differ from those attracted to traps baited with carbon dioxide or live hosts [9]. Additionally, sampling conditions such as the number of nights over which sampling occurs and weather conditions may also influence trapping success [10], with some species and traps potentially more affected than others. This variation in trapping outcome may not limit inference on the presence or absence of common species (e.g. the information used in global risk maps [11, 12]). However, it does limit inference based on comparing patterns of diversity or abundance across space or time [7]. Given that these comparisons are required to evaluate the ecological or anthropogenic drivers of mosquito populations and disease risk, choice of trapping methods presents challenges and opportunities for optimizing sampling efforts.

Previous studies have evaluated trapping methods in Europe, North America and South America [5–8, 13] while studies in southern Africa remain relatively limited. The mosquito fauna (Diptera: Culicidae) of southern Africa consists of over 216 species, many of which are endemic to the region [14]. Additional studies evaluating trapping methods are needed to evaluate if traps designed for other locations and species perform equally well in the species-rich communities in southern Africa. For example, a recent comparison of four traps in Germany found that the Biogents Sentinel trap (BG trap) collected the highest number of individuals in urban and snowmelt forest environments where *Culex* species predominate. In contrast, the Centers for Disease Control miniature light trap (CDC trap) collected the most mosquitoes in floodplain environments where *Aedes vexans* predominate, suggesting that the preferred trapping device may vary by habitat and community composition [7]. Qualitative evaluations in southern Africa suggest different traps likely collect different subsets of the mosquito community [15]. However, the two studies that systematically evaluated trapping methods in southern Africa focused only on house-based trapping in residential areas [16, 17].

An evaluation of trapping methods is also needed to influence the design of future entomological and pathogen surveillance efforts [18]. Such efforts could provide baseline information for public health interventions by identifying hotspots with a high abundance of vector species [19, 20]. There were more than 29,000 confirmed cases of malaria in southern Africa in 2017 (Botswana, Namibia, South Africa, Lesoto, Swaziland) [21]; key malaria vectors include members of the *Anopheles gambiae* complex (*An. arabiensis*, *An. gambiae*, *An. merus*) [22–24] and the *An. funestus* complex [25]. Mosquito-borne livestock and wildlife infections are also a concern, as outbreaks of West Nile virus, Rift Valley fever, Sindbis and Wesselsbron occur [26, 27]. Key vectors for these viral infections include, *Aedes caballus*, *Ae. circumluteolus*, *Ae. dentatus*, *Ae. juppi*, *Ae. mcintoshi*, *Ae. ochraceus*, *Culex pipiens*, *Cx. quinquefasciatus*, *Cx. univittatus* and *Cx. theileri* (based on suspected or known prime vectors in Africa, reviewed elsewhere [27]). Previous work has characterized the distribution [28], ecological drivers [29] and consequences for malaria risk [30] of key anopheline species. Understanding the distribution and drivers of non-Anophelinae mosquito species in southern Africa could facilitate informed management of a broader range of mosquito-borne disease or the development of vector control programmes that target multiple infections [15, 31].

In this study, we assessed four commonly-used adult mosquito trapping methods for their use in natural savanna ecosystems. Because surveillance programmes may have multiple aims (e.g. entomological surveys, pathogen surveillance or invasive/nuisance mosquito species control) we compared traps designed for a range of purposes with a range or attraction methods. We assessed the Centers for Disease Control miniature light trap (CDC) and a net trap due to their historic success for entomological surveys in a range of systems in southern Africa [9, 15, 16]. These traps use incandescent light + CO_2_ and CO_2_, respectively, to attract host-seeking vectors. We assessed the Biogents Sentinel-2 trap (BG) because of its worldwide success for both general entomological surveys [7] and targeted vector surveillance (e.g. *Ae. aegypti*, *Ae. albopictus*, *Cx. quinquefasciatus*, *Cx. pipiens* [10, 13]). It uses visual cues, CO_2_ and a lure to attract mosquitoes searching for hosts or a place to rest. We additionally compare the Biogents gravid *Aedes* trap (GAT) that is designed to collect gravid female mosquitoes, particularly container-breeding *Aedes* species, *Ae. aegypti* and *Ae. albopictus* [32, 33]. Gravid mosquitoes are more likely to be carrying pathogens, making the GAT trap potentially useful for pathogen surveillance or for joint pathogen and entomological surveillance efforts when used in combination with other traps.

We assessed these traps following two objectives. We compared estimates of abundance and community composition among all four traps (CDC, BG, GAT, net). We also evaluated traps designed for entomological surveillance (CDC, BG, net) by quantifying the importance of trapping method for inferring the regional distribution of mosquito abundance, mosquito community composition and the distribution of key disease vectors.

## Methods

### Study location

We sampled 16 sites within four regions of Kruger National Park (KNP), South Africa (Fig. 1; between 22°31’ and 25°31’S, 30°45’ and 32°00’E). We focus on KNP because it is a hotspot of mosquito diversity [15], it is in a region of southern Africa with regular outbreaks of mosquito-borne disease (e.g. [34, 35]), and it includes a sentinel site for mosquitoes and pathogen surveillance [36]). Sampling sites within KNP were chosen to cover the geographical extent of the park and to capture the variability in rainfall, geology and vegetation types within KNP [37]. This region experiences summer rainfall between November and April, and we selected our sampling to occur from March to April 2017, when mosquito populations are generally high. We sampled four out of the 22 management regions within the park: Malelane, Satara, Shingwedzi and Punda Maria. We additionally sampled in Skukuza during a pilot organizational and training week where traps were not directly compared. Within each region, we sampled at four sites, which were selected based on multiple criteria (Fig. 1). The primary selection criterion was to sample from water bodies that represented diverse types of wetlands (temporary ponds, permanent ponds, rivers). Additional criteria stipulated that the water bodies were at least 2 km away from one another to avoid sampling mosquitoes from adjacent water bodies and within 25 km from one another for sampling logistics. These distances are justified based on mean mosquito dispersal rates, which range from 35 m to 1.4 km depending on the species and habitat [38–41]. Although dispersal farther than 2 km is possible, it is uncommon [42] and we do not expect it to influence our results because only one pair of sites in Shingwedzi was located within 1 km of each other due limited surface water availability in the area.

**Fig. 1.**
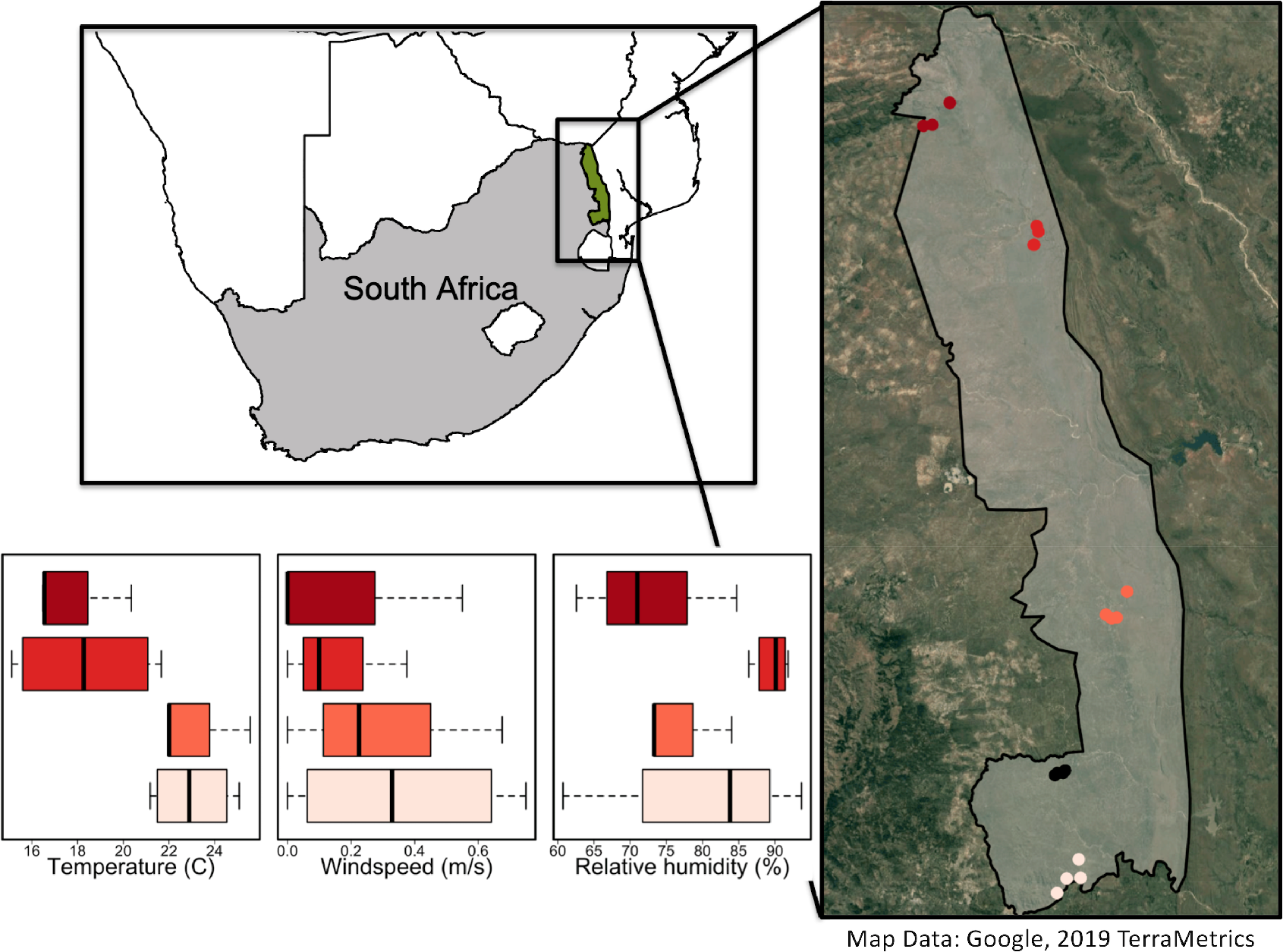
Map of the trapping sites within Kruger National Park (KNP), South Africa. Colors represent the regions where trapping occurred. From south to north, these include the Malelane, Skukuza (no weather data), Satara, Shingwedzi and Punda Maria sections. Each dot in the map represents a unique water body sampled. Regions of the park were characterized by distinct weather patterns (Additional file 1: Table S1). The base map uses Google, TerraMetrics imagery and was made using R with the GetMap function in the *RGoogleMaps* package [61]. The KNP boundary was provided by South African National Park’s Scientific Services

### Trapping and identification

We trapped at each site within a region for four consecutive nights and moved sequentially between regions each week in a random order (Skukuza: 20–23 March 2017; Malelane: 27–30 March 2017; Satara: 3–6 April 2017; Shingwedzi: 17–20 April 2017; Punda Maria: 24–27 April 2017). The four traps we used were the Centers for Disease Control miniature light trap with an incandescent light (Bioquip Inc, Rancho Dominguez, USA), a Biogents Sentinel 2 trap (Biogents AG, Regensburg, Germany), the Biogents gravid *Aedes* trap (Biogents AG) and the net trap. Although the net trap is not commercially available, it is easily and inexpensively made from netting and poles (see [15, 31]). For consistency in sampling, we set up the traps at a similar distance away from the water body (15–25 m) and approximately 30–50 m away from each other. Our aim was to estimate variation due to trap position, so we rotated the position of all traps after two nights of trapping. This study design allowed us to quantify how position within a site influences catch rates and compare if traps vary in their sensitivity to position and weather conditions (temperature, relative humidity and wind speed).

Our trap-use protocol was based on preliminary trapping work and expertise from mosquito surveillance in southern Africa [15, 31]. Specifically, we equipped the CDC trap, BG trap and net trap with a closed container containing 200–400 g of dry ice for a CO_2_ bait. To ensure the dry ice would last overnight, we reduced sublimation by wrapping the dry ice in multiple layers of newspaper and ensuring the container was closed except for 2 small holes. We placed the dry ice containers inside the BG traps, at the center of the net trap, and hanging with the CDC trap. Additionally, we equipped the BG traps with BG-Lures (Biogents AG). The GAT trap was baited with water following prior field trials suggesting water and hay-infusion bated GAT traps were equally successful [43]. We chose to use water to ensure consistently-baited traps across the study because although trap entry is not driven by chemical cues in the infusion [44, 45], oviposition behavior is highly variable across infusion types, infusion concentrations and fermentation period [33]. To set the trap, we placed a single piece of long-lasting insecticidal net inside the center of the trap, which was coated in 4.8% alphacypermethrin (Biogents AG). In addition to the spatial locations defined above, we hung the CDC light trap on a branch so it was approximately 1 m from the ground, and we placed the BG and GAT traps on the ground. We ensured the traps were placed close to the vegetation, but not directly under it, which has been shown to improve trapping success [10]. If available, we placed the net trap in slight shade and pulled the netting down to leave a gap of approximately 10 cm between the bottom of the net and the ground.

All traps were monitored twice a day at before-dusk (16:00–17:00 h) and dawn (6:00–7:00 h). The GAT trap was monitored during each visit but was collecting mosquitoes continuously because it does not require batteries. All other traps were set up before dusk and emptied again at dawn. The timing of this sample collection was used because the net trap requires clearing at dawn (mosquitoes may leave the trap after sunrise) and the CDC and BG traps require batteries, which we recharged during the day. This timing focuses our collection on species active between 16:00 h and the following 6:00 h, potentially missing species active just after dawn, the implications of which are discussed below. Due to rainfall, we excluded one trapping night at all sites in Satara and Punda Maria from all analyses.

Our sampling design (16 sites; 3–4 sampling nights each; 4 traps) resulted in 224 total trap-nights. Comparing traps at the same site resulted in 40 BG-CDC comparisons, 40 BG-GAT comparisons, 40 BG-net comparisons, 50 CDC-GAT comparisons and 50 CDC-net comparisons. The differences in sample size are due to both organization errors (11/224, or 5% of trapping nights) and losses due to wildlife interference (13/224, or 6% of trapping nights). Specifically, three BG traps were destroyed over the study by hyena, likely due to the BG-lure, which is specially designed to mimic human sent. Damage to traps from animals is an inherent feature of sampling in wildlife areas, although we took care to minimize the effects of our trapping. Because the consequences of these challenges have not previously been quantified and should be considered in future sample size calculations, we have provided detailed notes on trapping rates and trap-wildlife interactions in Additional file 1: Table S1. Our analyses are conservative in that they only compare data from the 220 successfully-collected, non-damaged trapping nights.

Directly after emptying the traps, we stored the mosquitoes on dry ice in the field and at −20 °C until identification. For identification of Anophelinae mosquitoes, we used identification literature from Gilles & Coutzee [46]. All anophelines were identified to species except for members of *An. funestus* complex and the *An. gambiae* complex, referred to herein as *An. gambiae* (*s.l.*), which require molecular methods for identification [38, 47]. For identification of the Culicinae, we used the key by Jupp [14]. Species were identified independently in duplicate by coauthors. Because species identification in the *Aedes vexans* complex (*Ae. vexans*, *Ae. hirsutus*, *Ae. fowleri*, *Ae. durbanensis*, *Ae. natronius*) and the *Aedes dentatus* complex (*Ae. dentatus*, *Ae. subdentatus Ae. pachyurus*, *Ae. bevisi*, *Ae. cumminsii*) were inconsistent at the species level, we aggregated them to the species complex level (Additional file 1, Table S2).

### Abiotic measurements

Temperature and rainfall within KNP follow a north-south gradient, with the highest values in the southwest [48]. We monitored temperature, relative humidity and wind speed using a Kestrel 3000 handheld weather meter (NK Inc., Boothwyn, PA, USA). We calculated the median temperature, relative humidity, and wind speed across sites within a region for each morning at the time of collection (Additional file 1: Table S3). All environmental variables were standardized for analysis, by centering and dividing by one standard deviation.

### Assessing how the number of traps and trapping nights influences mosquito richness

We assessed how observed species richness (the total number of unique species) saturates with sampling effort. We aggregated the data successively over 1, 2, 3 or 4 nights and calculated the cumulative proportion of species identified with an increasing proportion of nights. We also evaluated each trap’s ability to estimate richness by comparing richness estimated in pairs of non-damaged trap types at each site aggregated across all sampling nights using a Wilcoxon signed-rank test and a Bonferroni correction for multiple comparisons. This analysis accounts for trap losses by excluding comparisons from damaged traps.

### Assessing whether trap type influences estimates of mosquito abundance and inference on the regional patterns of abundance

To test for differences in abundance, we compared the number of mosquitoes collected per night between pairs of non-damaged traps. First, we quantified the relationship between trap type and abundance with a Wilcoxon signed-rank test and a Bonferroni correction for comparisons among the 4 traps. Then, we quantified the influence of trap type for inferences on the regional patterns of abundance using linear regressions with Poisson errors and a log-link function. The regression analyses assessed the relationship between a trap’s nightly mosquito counts with region of the park and weather conditions (wind speed, temperature, relative humidity). It also included random effect for site (Additional file 2: Table S4). We were additionally interested in the variance due to trap position within a site, estimated with a random effect for trap position. Although we were unable to estimate this parameter in the regression due to the number of damaged traps, our comparisons remain valid because the variation in counts due to trap position was smaller than the variation due to trap type or region. We fitted the regression model separately to data from the CDC trap, the net trap, the BG trap and data from aggregating abundance across all traps at a site. Because no individuals were collected in the GAT trap on most nights, we did not fit the model to data from the GAT trap alone. For each dataset, we conducted model selection using backward selection based on Akaike information criteria with a correction for small sample sizes (AICc) and select the model with the lowest AICc value. We fitted all regression models in R [49] using the *lme4* package [50].

### Assessing whether trap type influences estimates of mosquito community composition and inference on the regional patterns of mosquito communities

To assess if different traps provide different estimates of community composition, we first evaluated if certain species were particularly attracted to one trap over the other. We assessed species-specific trap bias by calculating the difference in the number of individuals for each species sampled between each pair of traps collected at a site over all nights of trapping. Because traps were paired at each site, we tested for differences between the traps using a Wilcoxon signed-rank test and a Bonferroni correction for multiple comparisons. We only compared the species-specific trap bias of common species, defined as being observed in the dataset more than three times. Because 23 common species were compared (k = 23), significant differences between traps occur when *P*-values are less than *P* = 0.05/23. We assessed trap bias for rare species visually.

To test for differences in community composition among traps, we used a non-parametric analysis of similarities analysis (ANOSIM), visualized potential differences with non-metric multidimensional scaling (NMDS) and quantified the influence of trap type for inferences on the regional patterns of community composition with hierarchical clustering. For all analyses, we calculated Bray-Curtis dissimilarity matrices based on the trap-specific (BG, CDC, net) abundances of all taxa within a site aggregated across sampling nights. The ANOSIM analysis is a non-parametric test for differences in mosquito communities among traps that compares the ranks of Bray-Curtis dissimilarity measures from samples collected from the same *vs* different traps [51]. To visualize this, we created an ordination of traps and sites in mosquito community space for each region of the park. Before all ordinations, we applied a Wisconsin transformation followed by a square root transformation to the species matrices, which standardizes by species maxima and reduces the influence of highly abundant taxa, respectively [52]. The ordinations converged on a stable two-dimensional solution, based on stress values. We conducted all community analyses in R using the *vegan* package [53].

### Describing regional patterns of disease vectors

We additionally describe how known prime vectors for West Nile virus (*Cx*. *pipiens*, *Cx. quinquefasciatus*, *Cx. theileri*, *Cx. univittatus*), Rift Valley fever (*Ae. dentatus*, *Ae. mcintoshi*, *Ae. ochraceus*), Sindbis (*Cx. univittatus*), and Wesselsbron (none found) are distributed across regions [22, 27]. Additional known prime vectors for these infections were not found in the study (*Ae*. *caballus*, *Ae. circumluteolus*, *Ae. juppi*). Chikungunya and dengue fever outbreaks are less common in South Africa [14], but we describe the distribution of their vector, *Ae. aegypti* [22], because additional known prime vectors in Africa were not found (*Ae. africanus*, *Ae. albopictus*, *Ae. cordellieri*, *Ae. furcifer*, *Ae. luteocephalus*, *Ae. neoafricanus*, *Ae. taylori*). We focus on the known prime vectors for a conservative description, but additional mosquito species are considered suspected vectors (reviewed in [27]). We used the numbers of *Cx. pipiens* complex to approximate the numbers of *Cx. pipiens* and *Cx. quinquefasciatus*, *Cx. univittatus* complex for *Cx. univittatus*, and *Ae. dentatus* complex for *Ae. dentatus.* Although this assumption may hide epidemiologically important variation, it is a valid approximation for these infections because multiple members of the complex are known or suspected vectors. However, we did not plot the distribution of malaria vectors because the most abundant species of *An. gambiae* (*s.l.*) in KNP, *An. quadriannulatus*, is not a malaria vector [15].

## Results

We collected 955 female mosquitoes, 946 (99.1%) of which were identified to the level of species or species complex. The most common species included members of the *Cx. univittatus* complex, *Ae. vexans* complex and *Cx. pipiens* complex. We also collected over 50 *An. gambiae* (*s.l.*), *An. pretoriensis* and *Cx. theileri* females. Mosquito data are provided in Additional file 3: Table S5.

### Mosquito communities can be characterized with the net and CDC traps after multiple trapping nights

Based on all traps together, mosquito community richness was sensitive to the number of sampling nights (Fig. 2; Additional file 4: Figure S1). Taken across all sites in Malelane, 89% (24/27) of the total number of unique species identified after four trapping nights were collected by night 2 and 96% (26/27) were collected by night 3. In Shingwedzi, 79% (19/24) of the species were collected by night 2 and 96% (23/24) by night 3. In Punda Maria, 53% (8/15) of the species were collected by night 2 and 87% (13/15) by night 3. These percentages overestimate the percent of richness captured because species accumulation curves suggest more than four nights of sampling are required to estimate total species richness (Fig. 2). The CDC and net trap together sampled all of the mosquito species captured (range across regions, 93–100%), while the BG and GAT trap captured far fewer species (Fig. 2). Estimates of richness in the BG trap were lower than in the net or CDC trap (Wilcoxon signed-rank test: BG *vs* net, *n* = 40, *V* = 136, *P* < 0.001; BG *vs* CDC, *n* = 40, *V* = 132.5, *P* < 0.001) but did not differ between the net or CDC trap (*n* = 50, *V* = 51, *P* = 0.940).

**Fig. 2.**
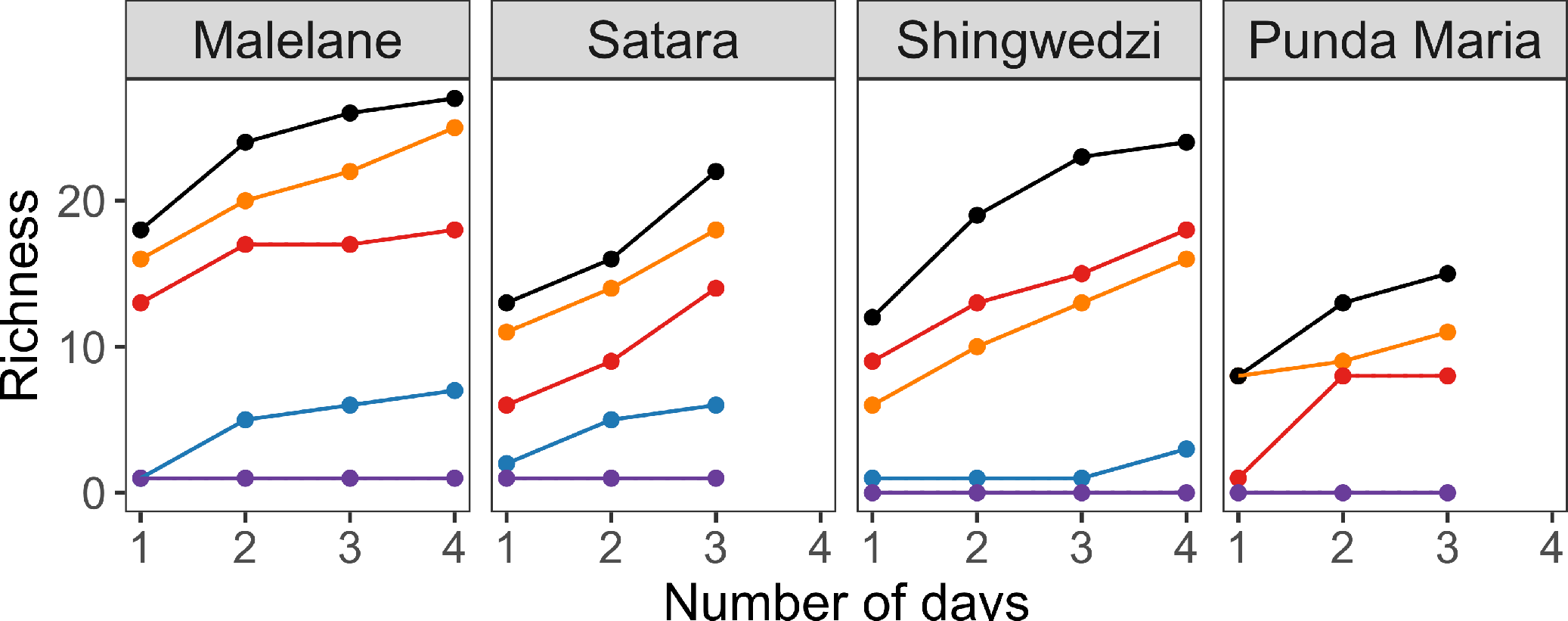
Richness (number of unique species) was sensitive to sampling effort and trap type. Richness values in each region were aggregated across sites. Data for each site within a region are provided in Additional file 1: Figure S1 and the number of traps represented in each region are specified in Additional file 1: Table S1. Sites within the Satara and Punda Maria region were only sampled for three nights due to rain

### Daily mosquito abundance estimates were comparable between the net and CDC traps but not the BG and GAT trap

Most individuals were collected in the CDC or the net trap, while the BG and GAT trap captured far fewer individuals (Fig. 3a). Together, the CDC and net trap sampled 96% of the individuals collected (range across regions, 94–99%; Additional file 1: Table S1). The net trap collected a mean of 8.6 females per night, the CDC trap collected a mean of 7.4, the BG trap collected a mean of 0.7, and the GAT trap collected a mean on 0.1 females (median: 5.5, 4, 0, 0). Paired by site, estimates of mosquito abundance based on the BG trap were lower than estimates from the net and CDC trap (BG *vs* CDC, *n* = 40, *V* = 22, *P* < 0.001; BG *vs* net, *n* = 40, *V* = 12, *P* < 0.001; Fig. 3b), while estimates based on the GAT trap were the lowest (BG *vs* GAT *n* = 40, *V* = 153, P < 0.001). However, estimates from the net and CDC trap did not significantly differ (CDC *vs* net, *n* = 50, *V* = 602.5, *P* = 0.12). When data from traps were aggregated across sites and nights within a region, trap choice continues to have a large influence on mosquito abundance (Additional file 1: Table S1).

**Fig. 3.**
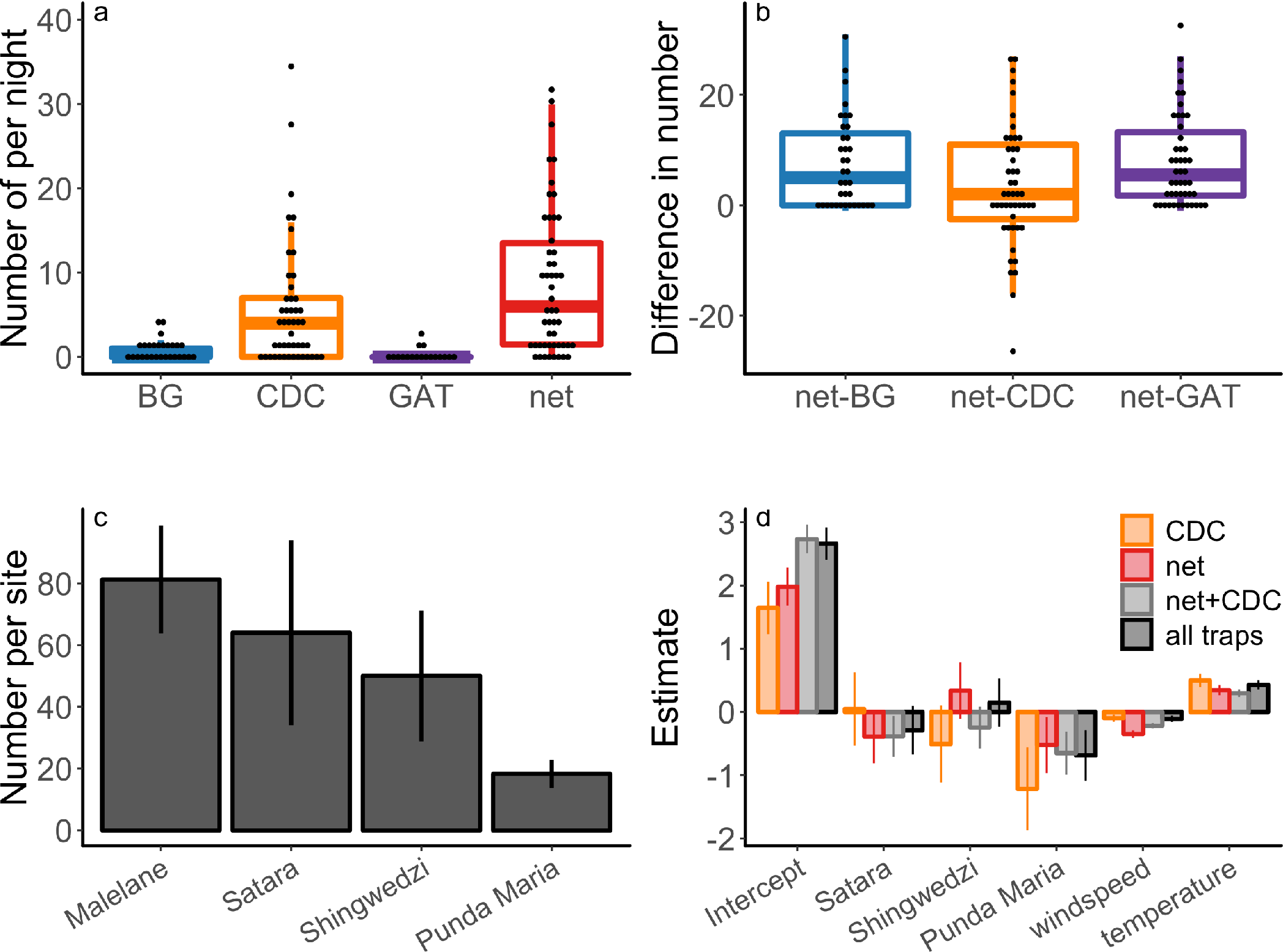
Mosquito abundance. **a** The number of mosquitoes captured by trap type; dots represent the number captured on each night at each site. **b** The difference in the number of mosquitoes sampled between trap types paired by sampling site. **c** The mean and standard error number of mosquitoes collected per site across region based on data aggregated across traps and sampling nights. **d** Regression parameter estimates and standard errors from statistical models characterizing the median number of mosquitoes sampled per night. Parameter values quantify the influence of sites relative to Malelane and weather conditions compared to the mean relative humidity (RH), temperature or wind speed. Colors indicate whether data used for model fitting was based on one trap or aggregated from multiple traps. The parameter estimates and hypothesis tests are defined in Additional file 1: Table S3

Regression analyses showed that trap choice also influences comparisons of abundance between regions. All traps identified similar trends, with the highest numbers captured in Malelane and the lowest numbers captured in Punda Maria (Fig. 3c, d). This spatial pattern was significantly different for models fit to the CDC data (Additional file 2: Table S4). All traps were influenced by weather conditions, particularly temperature and wind speed (Additional file 2: Table S4). We collected higher numbers in warm, low wind conditions (Fig. 3d). The net trap also collected higher numbers in low relative humidity conditions. Results based on counts aggregated from multiple traps were similar to results based on counts from either the CDC or net trap alone (Fig. 3c; Additional file 2: Table S4).

### Mosquito community composition was consistent between the net and CDC trap but not the BG trap

Community composition was similar between the net and CDC trap, but not for the BG trap (ANOSIM, overall: *R* = 0.126, *P* = 0.04; pairwise: net *vs* CDC, *P* = 0.894; net *vs* BG, *P* = 0.023; CDC *vs* BG, *P* = 0.009). NMDS ordinations of traps in species space reflect this relationship although some variation across regions of the park does exist (Additional file 4: Figure S2, Table S6). However, we did not find evidence for species-specific bias between any of the traps, suggesting that differences in community composition in the BG trap are driven by the relatively lower abundance collected in the trap (Fig. 4). For common species collected in the net *vs* CDC trap, the number of individuals collected was not significantly different between the traps (Fig. 4a; no hypothesis tests for individual species were significant). Rare species also did not show trap biases and include *Ae. aerarius*, *Ae. metallicus*, *Ae. unidentatus*, *An. maculipalpis*, *An. ziemanni*, *Cx. antennatus*, *Cx. bitaenorhynchus*, *Cx. nebulosus*, *Lutzia tigripes*, *Mansonia africana* and *Uranotaenia balfouri* (Additional file 4: Figure S3). For comparisons with the BG trap, the net and CDC traps both collected higher numbers of individuals, but there were no species or genus shifts driving this pattern (Fig. 4b, c; Additional file 4: Figure S4).

**Fig. 4.**
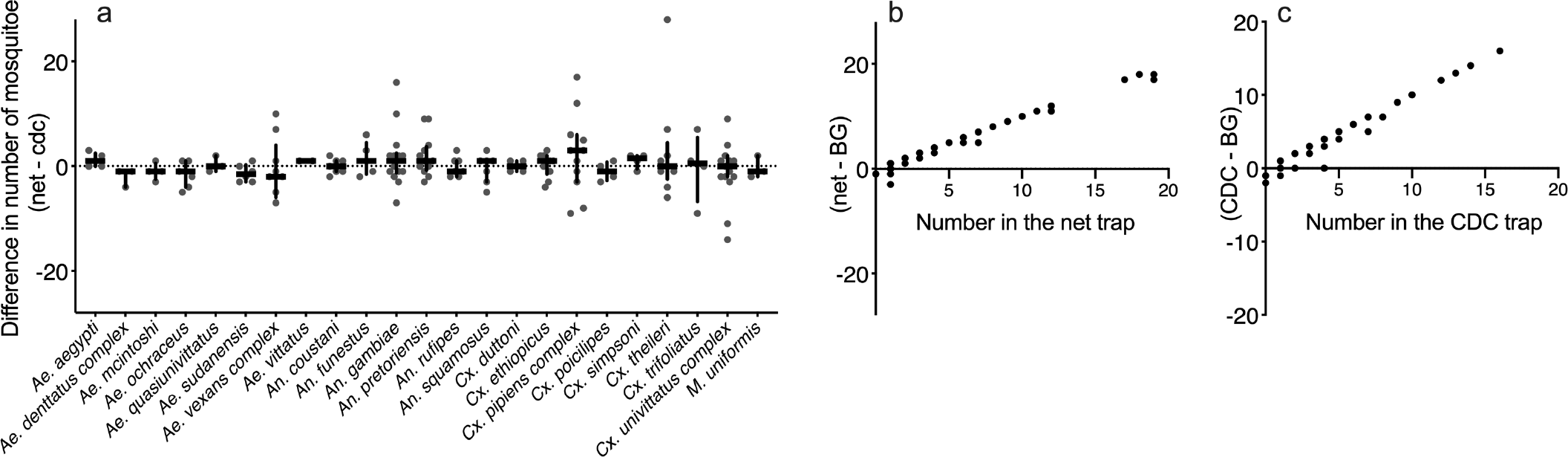
There were no species-specific differences between the net and CDC trap (**a**), while species-specific differences between the net and BG trap (**b**) and the CDC trap and BG trap (**c**) were driven by overall abundance in the net or CDC trap. Dots represent the difference in the number of mosquitoes collected between pairs of traps based on the total number of mosquitoes sampled across nights at each site. **a** Lines represent the median and interquartile range of the data. Data displayed do not include sites where no individuals of a given species were collected in either trap, but results remain consistent regardless of whether these sites are or are not included. No hypothesis test for the individual species was significant (Wilcoxon signed-rank test, *P* = 0.181, 0.174, 0.345, 0.143, 1, 0.134, 0.476, 0.346, 1, 0.498, 0.360, 0.152, 0.796, 0.931, 1, 0.719, 0.372, 0.423, 0.265, 0.725, 1, 0.850, 1). See Additional file 1: Figure S3 for species-specific comparisons with the BG trap and Additional file 1: Table S5 for a summary table. *Abbreviations*: Punda, Punda Maria; wind, wind speed; temp, temperature

Although trap choice influences estimates of community composition (e.g. community richness, ordinations), hierarchical cluster analysis suggests that may be less important for comparisons between regions (Fig. 5; Additional file 4: Figure S5). Regional patterns in mosquito communities were consistent across trap types, with samples from Malelane and Satara being more similar than samples from Shingwedzi. Mosquito communities estimated from the CDC and net trap were clustered by region (Additional file 4: Figure S5), indicating that communities within a region were more closely related to each other than communities between regions regardless of the net or CDC trap. In contrast, samples from the BG trap were clustered separately from the samples from the net and CDC trap (Fig. 5). We therefore describe the regional patterns of disease vectors based on data from all traps together (Fig. 6). Rift Valley fever vectors were most common in Satara while West Nile virus and Sindbis vectors were more common in Shingwedzi and Punda Maria. We did not find known vectors for Wesselsbron (*Ae. caballus*, *Ae. circumluteolus*) within the park.

**Fig. 5.**
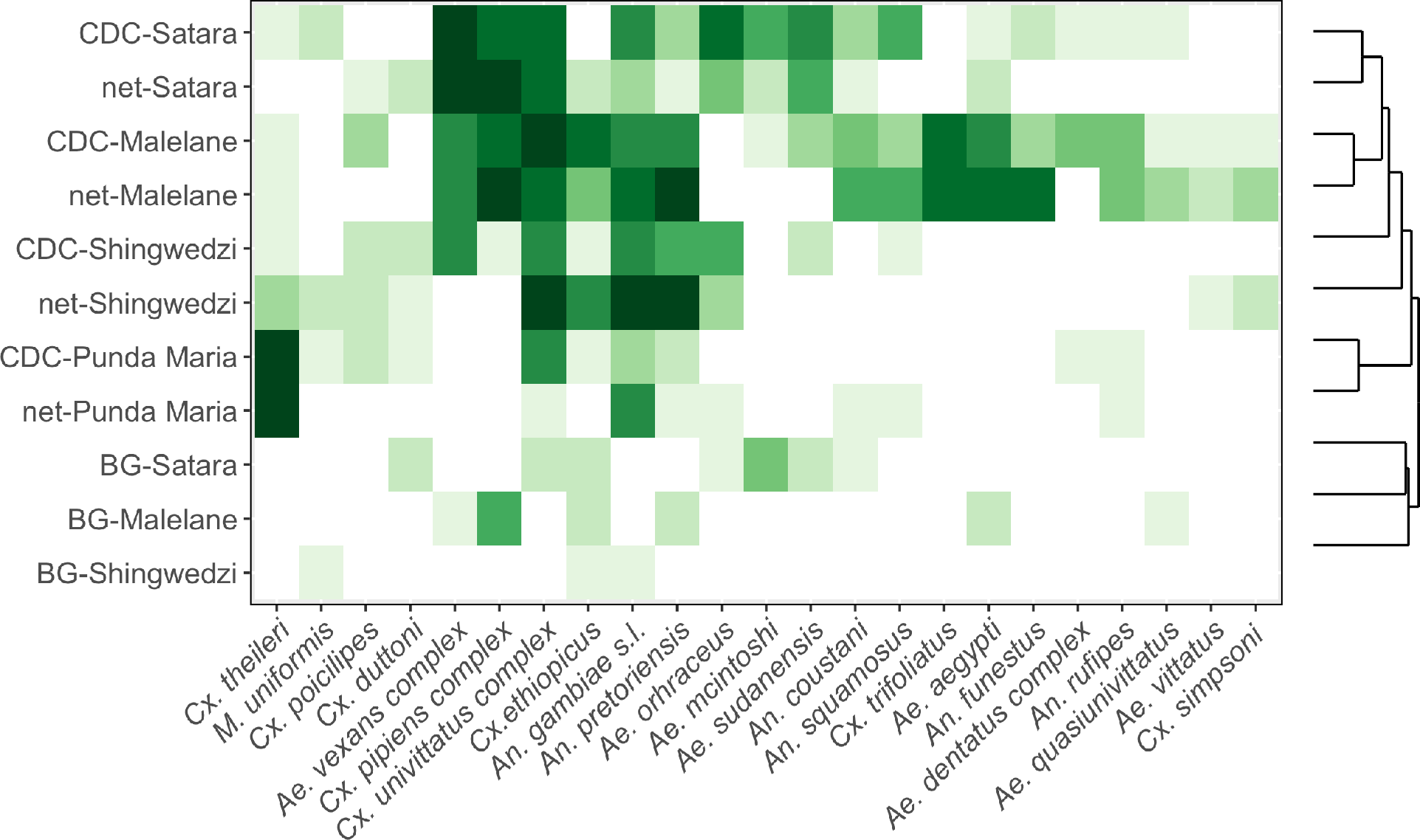
Communities collected in the net and CDC trap were clustered by region. Traps and regions are ordered according to the tree produced by clustering (Additional file 1: Figure S5). Colors represent species abundance, with color bins defining the 30th to 90th percentile in increments of 10

**Fig. 6.**
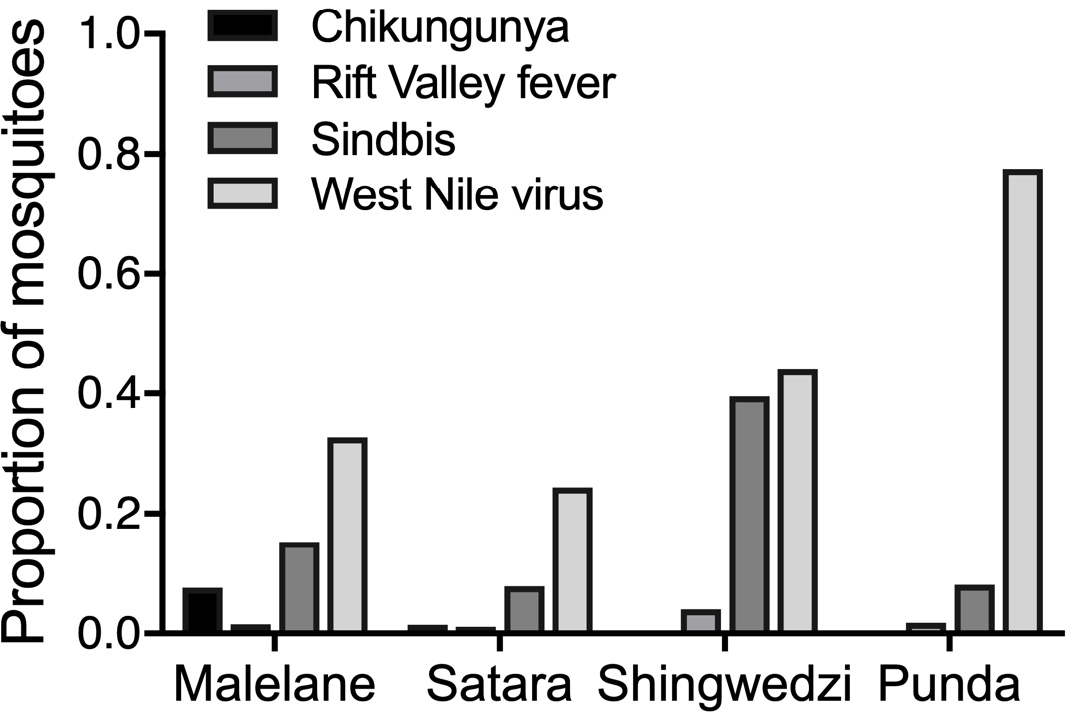
The proportion of mosquitoes identified as primary vectors in each region. *Abbreviation*: Punda, Punda Maria

## Discussion

Mosquito trapping is used for disease surveillance, biodiversity surveys and nuisance-reduction. In light of these multiple, non-exclusive aims, this study compared traps based on four potential goals: collecting large numbers of mosquitoes; estimating mosquito community composition, including invasive, vector or rare species; characterizing the spatial patterns of abundance; and characterizing the spatial patterns of community composition. We expected trade-offs among these aims, for example with traps specializing on one vector to be less successful in estimating community richness (and *vice versa*, e.g. [54]). In contrast to this expectation, our results indicated that the net and CDC trap consistently performed best across multiple outcomes.

The CDC and net trap collected higher abundances and more unique species compared to the BG and GAT traps. These differences are based on comparisons in 40 or more trapping nights from 16 sites where both traps were deployed (Additional file 1: Table S1). This result is different from trap comparison studies in Europe [7], the USA (BG [13], GAT [55]) and South America (BG [56], GAT [57]), where both the BG and GAT traps have been shown to perform well. One reason for their relatively low success within KNP could be the diversity and types of species present within the park. The GAT trap has been designed to capture container-breeding species, such as *Ae. aegypti* [57], and the BG trap performs well in sampling *Ae. aegypti*, *Ae. albopictus* and *Cx. pipiens* [7, 58]. Although *Ae. aegypti* was present in our study and previous studies [15], they were relatively rare (Additional file 1: Table S2). The BG trap’s sampling efficiency for the two most-common species complexes in our dataset, members of the *Ae. vexans* complex and *Cx. univitatus* complex, is either low [7] for the *Ae. vexans* complex or previously unknown for the *Cx. univitatus* complex. Future work might find improved catch rates for the BG trap by modifying the type of lure used or extending the sampling period to 9:30 h to catch *Aedes* species that bite at dusk and dawn (peak *Ae. aegypti* activities between 15:30–19:30 h and 5:30–9:30 h; [59]). An alternative reason for the low success of the BG trap could be that it is negatively influenced by the presence of alternative refugia and oviposition sites [60]. Habitat heterogeneity is likely to be higher within KNP compared to other, primarily urban or suburban environments where mosquito traps have been evaluated ([13, 55, 56] but see [7]). Additional comparisons in urban environments in southern Africa are needed to distinguish these two hypotheses.

By providing a detailed comparison between the net and CDC traps, our results suggest that these traps provide similar estimates of community richness and community diversity. A key aim of this work was to evaluate how choice of trap influences spatial estimates of mosquito abundance and mosquito community composition. As a result, our sites were selected to sample a diversity of breeding habitats, some of which were known to have high numbers of mosquitoes and others which we anticipated to have lower numbers. This approach is a better approximation to how the traps would be deployed for surveillance compared to targeting only sites with high numbers of mosquitoes. Despite this, the CDC trap collected 7.4 females per night while the net trap collected 8.6 females per night, similar to previous collection efforts in KNP that report aggregated data, with a median of 31 (range 17–116) females across three net traps and a median of 19 (range 19–33) in three CDC traps [15]. Based on this data, the consistent patterns of community composition and largely consistent patterns of abundance between traps suggest that comparisons between studies using these two methods are possible.

This work sets the stage for follow-up studies comparing mosquito abundance and community composition across space or time. We recommend that such a follow-up study should use the CDC or net trap across multiple nights or sites within a region. More traps and sampling will be needed to characterize spatial patterns of abundance because we observed significant variation among sites and relatively low abundances per trap. The choice between the CDC and net trap should also consider other features of the traps, such as ease of use and specimen quality. For morphological identification, the net trap has the advantage that mosquito specimens can be collected with minimal damage, which makes them easier to identify [14]. It also has no motorized or battery-powered parts, making it difficult to break and straightforward to mend. Therefore, studies requiring precise species identification such as biodiversity assessments or studies conducted in remote locations may prefer the net trap. However, for sampling large numbers of sites or sites in remote locations, the CDC trap has an advantage because the timing of when traps need to be cleared is more flexible compared to the net trap, which has to be cleared at sunrise. These practical considerations may mean that the CDC trap is better suited for large, comparative studies.

## Conclusions

After assessing four different mosquito trapping methods in a natural savanna ecosystem, we recommend the net trap, the CDC trap or their combined use for outdoor mosquito surveillance in southern Africa. These traps performed well based on four evaluation criteria: collecting large numbers of mosquitoes; estimating mosquito community composition, including vector or rare species; characterizing the spatial patterns of abundance; and characterizing the spatial patterns of community composition. This suggests they are appropriate for both biodiversity surveys and vector surveillance. As such, this study provides a valuable proof-of-principle for characterizing the spatial patterns of non-vectors as well as vectors for multiple diseases.

## Additional files

**Additional file 1: Table S1.** The number of females collected by trap and site. The mosquito column (mosq) indicates the number of females collected; the traps column of trapping nights represented. **Table S2**. The number of females collected of each species by region of the park. **Table S3**. Summary of weather conditions sampled within each region of the park. Numbers are displayed as median and range in parentheses.

**Additional file 2: Table S4**. Model parameters, estimates, standard error (SE) and hypothesis tests for the Poisson regression analyses in Fig. 3.

**Additional file 3: Table S5.** Data on the number of species collected.

**Additional file 4: Table S6**. Descriptive results comparing species-specific shifts in mosquito communities collected in the net and CDC trap. Species more commonly collected in a trap are listed if 5 more were collected in that trap after all sampling days at the site. **Figure S1**. The apparent richness (number of unique species) and diversity for sites within each region. **Figure S2**. Non-metric multidimensional scaling ordinations of trap differences in mosquito communities in Malelane, Satara, Shingwedzi and Punda Maria. **Figure S3**. Species-specific trap preferences for the net *vs* CDC trap difference based on rare species not displayed in Fig. 3. Dots represent the difference in the number of mosquitoes collected in the net *vs* the CDC trap based on the total number of mosquitoes sampled across nights at each site. **Figure S4**. The net trap and the CDC trap caught higher numbers of mosquitoes (Fig. 3) and this pattern was not driven by any species or genus-specific trap bias (left figures) but by variation in the total number of the species collected (right figures). **Figure S5**. Dendrogram of species composition based on Bray-Curtis dissimilarity and the hierarchical clustering algorithm.

## Abbreviations

AICc: Akaike information criteria with a correction for small sample sizes
ANOVA: analysis of variance
BG: Biogents sentinel trap
CDC: Centers for Disease Control miniature light trap
GAT: Biogents gravid *Aedes* trap
KNP: Kruger National Park
SE: standard error.

## Acknowledgements

We thank South African National Parks (SANParks) for their support in conducting this study in Kruger and providing us with mosquito specimens to help with identification. We thank Purvance H. Shikwambana and a great field team: Vicky Beckers, Nina Haver, Louie Krol, Skhumbuza Mdletshe, Karabo Moloi, Nondumiso Myataza and Gijs van Nes. We thank Dr Alan Kemp, Professor A. Paulo Gouveia de Almeida and Dr Anthony Cornel for their assistance with identification.

## Declarations

### Ethics approval and consent to participate

Not applicable.

### Consent for publication

Not applicable.

### Availability of data and materials

The datasets supporting the conclusions of this article are available in Additional file 3 and the free online mosquito database, VectorMap (http://vectormap.si.edu).

### Authors’ contributions

EEG, BRB and MS conceived and designed the analysis; EEG, BRB, DG and MS collected the data; EEG, MMG and MS conducted identification; EEG, PMB and MS performed analyses; all authors wrote the paper. All authors read and approved the final manuscript.

