## Additional File for "A comparative assessment of adult mosquito trapping methods to estimate spatial patterns of abundance and community composition in southern Africa"

### Additional File 1

#### Additional descriptive data tables

*Mosquito data*

The data capture 200 total trap-nights and resulted in 40 BG-CDC comparisons, 40 BG-GAT comparisons, 40 BG-net comparisons, 50 CDC-GAT comparisons, and 50 CDC-net comparisons. Below, we provide summary statistics for the mosquito data aggregated across 3-4 sampling nights at each site (Table S1) and aggregated across sampling nights for each species (Table S3). Sites vary in trapping effort due to rainfall in Satara and Punda Maria as well as animal interference. This aggregated data is displayed for summary purposes, as analyses of mosquito abundance compare the number of mosquitoes collected per night between functioning traps.

*Weather data*

Regions of the park were characterized by distinct weather patterns, as temperature was significantly different between regions (ANOVA: temperature, $F_{(3, 10)} = 4.88$, $P<0.024$; wind speed, $F_{(3, 10)} = 0.41$, $P = 0.752$; relative humidity, $F_{(3, 10)} = 1.97$, $P = 0.183$).

**Table S1.** The number of females collected by trap and site. The mosquito column (mosq) indicates the number of females collected; the traps column of trapping nights represented.

| Trap | BG | | CDC | | GAT | | net | | Notes |
|---|---|---|---|---|---|---|---|---|---|
| | mosq | traps | mosq | traps | mosq | traps | mosq | traps | |
| Malelane Total | 14 | 14 | 150 | 16 | 4 | 15 | 159 | 15 | (total - across 4 nights) |
| Malelane 1 | 5 | 4 | 79 | 4 | 0 | 4 | 19 | 4 | |
| Malelane 2 | 4 | 3 | 38 | 4 | 0 | 4 | 72 | 4 | |
| Malelane 3 | 1 | 3 | 15 | 4 | 0 | 3 | 20 | 4 | |
| Malelane 4 | 4 | 4 | 18 | 4 | 4 | 4 | 48 | 3[a] | a. Net not set-up, elephants |
| Satara Total | 12 | 9 | 175 | 11 | 1 | 12 | 80 | 11 | (total - across 3 nights) |
| Satara 1 | 9 | 3 | 26 | 3 | 0 | 3 | 24 | 3 | |
| Satara 2 | 1 | 2 | 15 | 3 | 1 | 3 | 23 | 3 | |
| Satara 3 | 0 | 1[b] | 128 | 3 | 0 | 3 | 23 | 2 | b. Two BG traps destroyed, hyena |
| Satara 4 | 0 | 3 | 6 | 2[c] | 0 | 3 | 10 | 3 | c. CDC pulled down, baboons |
| Shingwedzi Total | 3 | 10 | 51 | 15 | 0 | 15 | 146 | 15 | (total - across 4 nights) |
| Shingwedzi 1 | 2 | 3 | 0 | 3 | 0 | 3 | 21 | 3 | |
| Shingwedzi 2 | 0 | 2[d] | 11 | 4 | 0 | 4 | 9 | 4 | d. BG trap not deployed, previous damage |
| Shingwedzi 3 | 1 | 3 | 17 | 4 | 0 | 4 | 93 | 4 | |
| Shingwedzi 4 | 0 | 2[c,d] | 23 | 4 | 0 | 4 | 23 | 4 | d. BG trap not deployed, previous damage<br>b. One BG trap destroyed, hyena |
| Punda Maria Total | 1 | 8 | 18 | 11 | 0 | 12 | 62 | 11 | (total - across 3 nights) |
| Punda Maria 1 | 1 | 3 | 1 | 3 | 0 | 3 | 12 | 3 | |
| Punda Maria 2 | 0 | 2[d] | 0 | 2 | 0 | 3 | 31 | 3 | d. BG trap not deployed, previous damage |
| Punda Maria 3 | 0 | 2[d] | 7 | 3 | 0 | 3 | 11 | 3 | d. BG trap not deployed, previous damage |
| Punda Maria 4 | 0 | 1[d] | 10 | 3 | 0 | 3 | 8 | 2[e] | d. BG trap not deployed, previous damage<br>e. tent trap fell over, unknown animal |
| **Overall Total** | 30 | 41 | 349 | 53 | 5 | 54 | 447 | 52 | |

**Table S2.** The number of females collected of each species by region of the park.

| Species | Malelane | Skukuza | Satara | Shingwedzi | Punda Maria | Total |
|---|---|---|---|---|---|---|
| *Aedes aegypti* | 25 | 0 | 4 | 0 | 0 | 29 |
| *Aedes aerarius* | 2 | 0 | 0 | 0 | 0 | 2 |
| *Aedes dentatus* complex | 4 | 0 | 1 | 0 | 1 | 6 |
| *Aedes mcintoshi* | 1 | 0 | 11 | 0 | 0 | 12 |
| *Aedes metallicus* | 0 | 0 | 1 | 1 | 0 | 2 |
| *Aedes ochraceus* | 0 | 0 | 19 | 8 | 1 | 28 |
| *Aedes quasiunivittatus* | 5 | 0 | 1 | 0 | 0 | 6 |
| *Aedes sudanensis* | 3 | 1 | 15 | 2 | 0 | 21 |
| *Aedes unidentatus* | 3 | 0 | 0 | 0 | 0 | 3 |
| *Aedes vexans* complex | 18 | 1 | 120 | 9 | 0 | 148 |
| *Aedes vittatus* | 3 | 0 | 0 | 1 | 0 | 4 |
| *Anopheles coustani* | 9 | 4 | 5 | 0 | 1 | 19 |
| *Anopheles funestus* | 15 | 0 | 2 | 0 | 1 | 18 |
| *Anopheles gambiae s.l.* | 23 | 0 | 12 | 32 | 9 | 76 |
| *Anopheles maculipalpis* | 0 | 0 | 0 | 3 | 0 | 3 |
| *Anopheles pretoriensis* | 28 | 1 | 4 | 24 | 3 | 60 |
| *Anopheles rufipes* | 8 | 0 | 1 | 0 | 2 | 11 |
| *Anopheles squamosus* | 8 | 0 | 5 | 1 | 1 | 15 |
| *Anopheles ziemanni* | 0 | 0 | 0 | 1 | 0 | 1 |
| *Culex antennatus* | 0 | 0 | 0 | 1 | 0 | 1 |
| *Culex bitaenorhynchus* | 0 | 0 | 0 | 1 | 0 | 1 |
| *Culex duttoni* | 0 | 0 | 4 | 3 | 1 | 8 |
| *Culex ethiopicus* | 17 | 5 | 2 | 11 | 1 | 36 |
| *Culex nebulosus* | 0 | 0 | 0 | 1 | 0 | 1 |
| *Culex pipiens* complex | 55 | 0 | 45 | 5 | 0 | 105 |
| *Culex poicilipes* | 3 | 11 | 1 | 4 | 2 | 21 |
| *Culex simpsoni* | 4 | 0 | 0 | 2 | 0 | 6 |
| *Culex theileri* | 2 | 0 | 1 | 4 | 77 | 84 |
| *Culex trifoliatus* | 33 | 2 | 0 | 0 | 0 | 35 |
| *Culex univittatus* complex | 50 | 10 | 22 | 79 | 9 | 170 |
| *Lutzia tigripes* | 0 | 0 | 0 | 0 | 1 | 1 |
| *Mansonia africana* | 1 | 3 | 0 | 2 | 0 | 6 |
| *Mansonia uniformis* | 0 | 0 | 2 | 3 | 1 | 6 |
| *Uranotaenia balfouri* | 0 | 0 | 0 | 1 | 0 | 1 |
| 3 unidentified *Anopheles* species | 2/1/3 | 0/0/0 | 0/0/0 | 0/0/0 | 0/0/0 | 2/1/3 |
| Unidentified *Culex* species | 1 | 0 | 0 | 0 | 0 | 1 |
| Unidentified *Mimomyia* species | 0 | 0 | 1 | 0 | 0 | 1 |
| Unidentified Genus | 0 | 0 | 0 | 1 | 0 | 1 |

**Table S3.** Summary of weather conditions sampled within each region of the park. Numbers are displayed as median and range in parentheses.

| | temperature | wind speed | relative humidity |
|---|---|---|---|
| Malelane | 22.9 (21.2-25.1) | 0.33 (0-0.75) | 83.8 (60.7-93.7) |
| Satara | 22.0 (21.7-25.6) | 0.23 (0-0.68) | 73.3 (73.3-84.0) |
| Shingwedzi | 18.3 (15.1-21.7) | 0.10 (0-0.38) | 90.1 (86.4-91.8) |
| Punda Maria | 16.6 (16.5-20.4) | 0.00 (0-0.55) | 71.0 (62.6-84.7) |

### Additional File 2

#### Additional methods and results for the regression analysis

All models consider the counts of mosquito from each night $i$ with the response variable, $\mu_i$. Explanatory variables include a factor variable for the effect of the region that trapping occurred in ($region_k$) and continuous variables for the effect of wind speed, temperature and relative humidity ($windspeed_i$, $temperature_i, RH_i$). We additionally included a site-specific intercept, $\propto_{j[i]}$, with subscript notation to indicate which of the $j$ sites were sampled in night *i*. This results in the following regression model,

$$\ln(\mu_i) = \propto_{j[i]} + \beta_{1,k} region_k + \beta_2 windspeed_i + \beta_3 temperature_i + \beta_4 RH_i + \varepsilon_i$$

where $\varepsilon_i$ is the Poisson distributed random error.

Following model selection, we evaluate $\beta_{1,k}$ to determine how the average nightly counts of mosquitoes in region $k$ differed from counts in Malelane, and we evaluate $\beta_2, \beta_3, \beta_4$ to determine how the average counts of mosquitoes vary with wind speed, temperature, and relative humidity, respectively. If the relative values of $\beta_{1,k}$ are consistent among traps (e.g. $\beta_{1,1} > \beta_{1,2} > \beta_{1,3}$) for models fit to data from different traps, then we conclude that trap choice does not influence spatial comparisons among regions. We were additionally interested in the variance components, as high variances indicate that counts vary across sites. We did not estimate variation due to trap position due to the number of damaged traps. However, regression models fit with the additional random effect to data from a subset of sites with full trap data resulted in similar estimates.

**Table S4.** Model parameters, estimates, standard error (SE) and hypothesis tests for the Poisson regression analyses in Fig. 3.

| **Model** | **Estimate** | **SE** | **Z value** | **P** |
|---|---|---|---|---|
| **BG data** (n = 41; variance across sites = 0.062) | | | | |
| $\beta_{1,1}$ – Satara vs. Malelane | 0.280 | 0.449 | 0.623 | 0.533 |
| $\beta_{1,2}$ – Shingwedzi vs. Malelane | -0.073 | 0.828 | -0.089 | 0.930 |
| $\beta_{1,3}$ – Punda Maria vs. Malelane | -0.935 | 1.183 | -0.791 | 0.429 |
| $\beta_3$ – temperature | 0.700 | 0.330 | 2.117 | 0.034 |
| **CDC data** (n = 52; variance across sites = 0.621) | | | | |
| $\beta_{1,1}$ – Satara vs. Malelane | 0.049 | 0.579 | 0.085 | 0.933 |
| $\beta_{1,2}$ – Shingwedzi vs. Malelane | -0.508 | 0.613 | -0.829 | 0.407 |
| $\beta_{1,3}$ – Punda Maria vs. Malelane | -1.215 | 0.657 | -1.850 | 0.064 |
| $\beta_2$ – wind speed | -0.098 | 0.059 | -1.653 | 0.098 |
| $\beta_3$ – temperature | 0.498 | 0.102 | 4.864 | <0.001 |
| **Net data** (n = 52; variance across sites = 0.326) | | | | |
| $\beta_{1,1}$ – Satara vs. Malelane | -0.388 | 0.424 | -0.916 | 0.359 |
| $\beta_{1,2}$ – Shingwedzi vs. Malelane | 0.338 | 0.445 | 0.759 | 0.448 |
| $\beta_{1,3}$ – Punda Maria vs. Malelane | -0.522 | 0.443 | -1.179 | 0.238 |
| $\beta_2$ – wind speed | -0.348 | 0.063 | -5.545 | <0.001 |
| $\beta_3$ – temperature | 0.344 | 0.079 | 4.336 | <0.001 |
| $\beta_4$ – relative humidity | -0.271 | 0.071 | -3.823 | 0.001 |
| **Net + CDC data** (n = 50, variance across sites = 0.181) | | | | |
| $\beta_{1,1}$ – Satara vs. Malelane | -0.385 | 0.330 | -0.742 | 0.458 |
| $\beta_{1,2}$ – Shingwedzi vs. Malelane | -0.245 | 0.330 | -0.724 | 0.458 |
| $\beta_{1,3}$ – Punda Maria vs. Malelane | -0.651 | 0.340 | -1.916 | 0.055 |
| $\beta_2$ – wind speed | -0.217 | 0.044 | -4.909 | <0.001 |
| $\beta_3$ – temperature | 0.298 | 0.062 | 4.782 | <0.001 |
| **All data combined** (n = 38, variance across sites = 0.222) | | | | |
| $\beta_{1,1}$ – Satara vs. Malelane | -0.288 | 0.382 | 0.752 | 0.452 |
| $\beta_{1,2}$ – Shingwedzi vs. Malelane | -0.148 | 0.382 | -0.387 | 0.699 |
| $\beta_{1,3}$ – Punda Maria vs. Malelane | -0.688 | 0.404 | -1.704 | 0.088 |
| $\beta_2$ – wind speed | -0.107 | 0.051 | -2.095 | 0.036 |
| $\beta_3$ – temperature | 0.432 | 0.076 | 5.702 | <0.001 |

### Additional File 4

#### Additional results

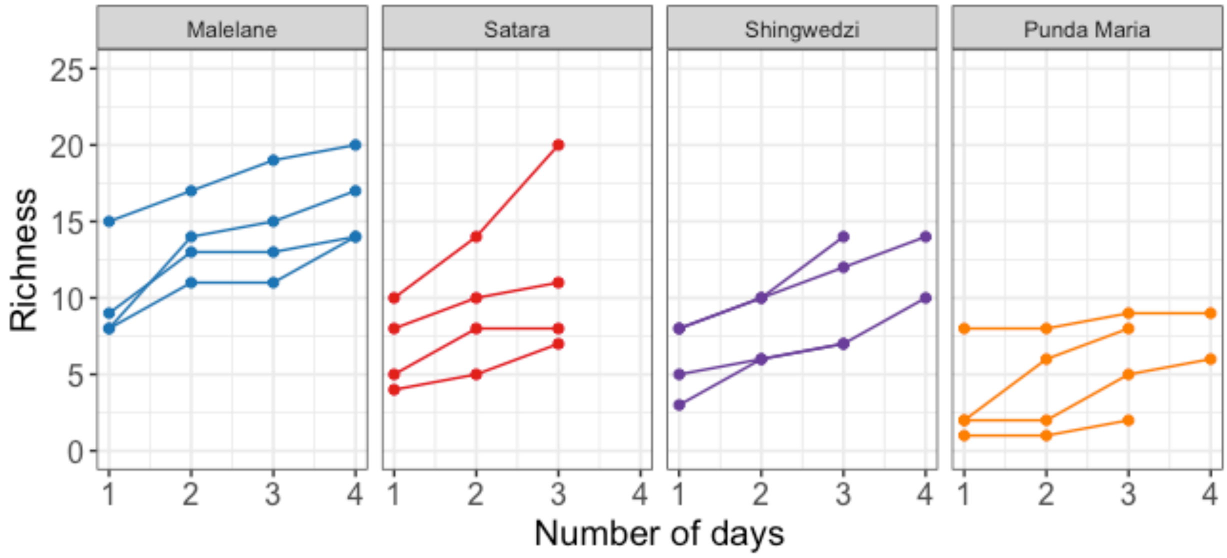

**Figure S1.** The apparent richness (number of unique species) and diversity for sites within each region.

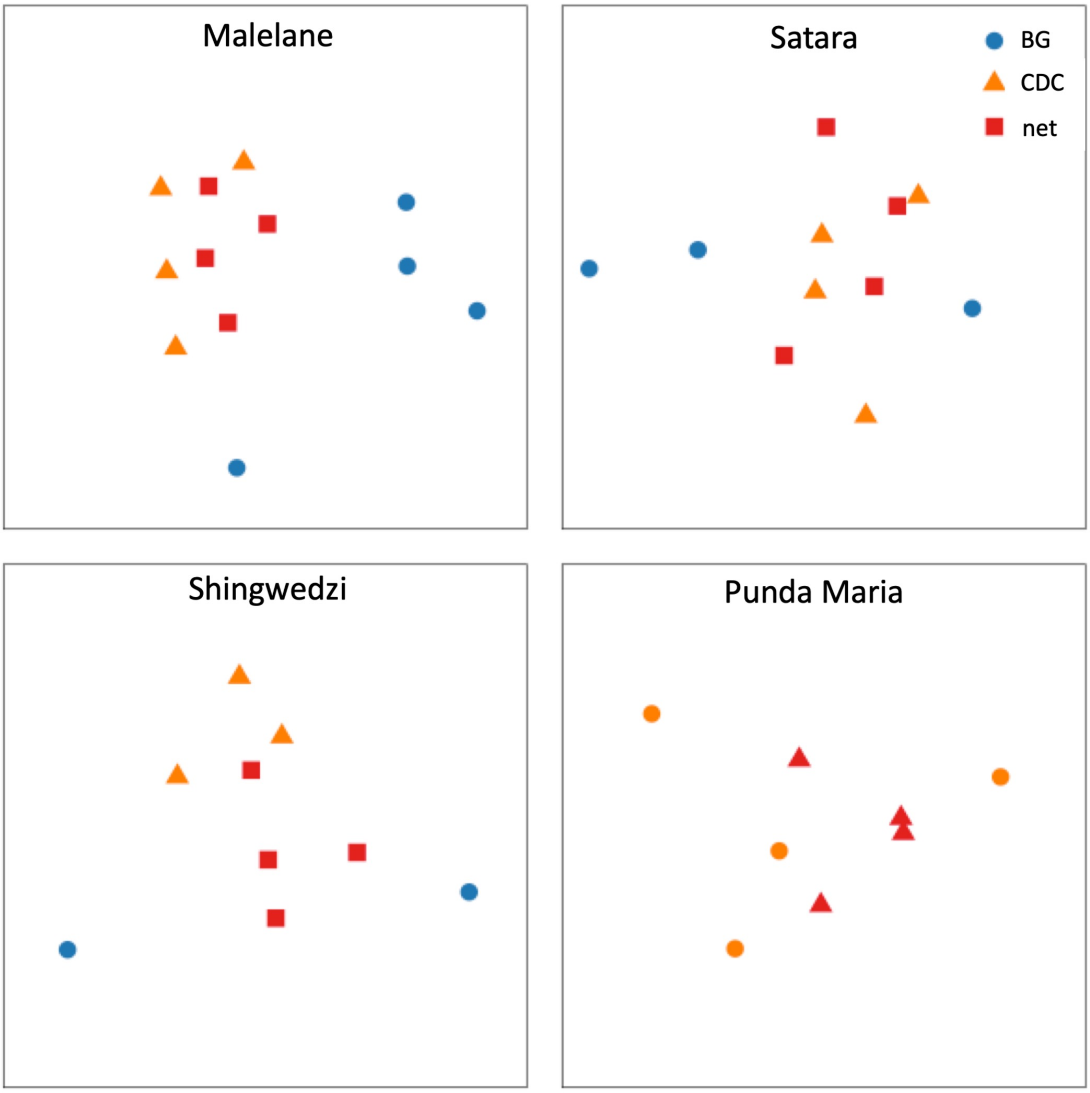

**Figure S2.** Non-metric multidimensional scaling ordinations of trap differences in mosquito communities in Malelane, Satara, Shingwedzi and Punda Maria.

**Table S6.** Descriptive results comparing species-specific shifts in mosquito communities collected in the net and CDC trap. Species more commonly collected in a trap are listed if 5 more were collected in that trap after all sampling days at the site.

| Site | Species common to the CDC trap | Species common to the net trap |
|---|---|---|
| Malelane 1 | *C. univittatus* complex*; C. trifoliatus; C. pipiens* complex*; Ae. vexans* complex | - |
| Malelane 2 | *C. univittatus* complex | *An. gambiae s.l.; An. pretoriensis; C. trifoliatus; Ae. vexans* complex |
| Malelane 3 | - | - |
| Malelane 4 | - | *C. pipiens* complex |
| Satara 1 | *C. pipiens* complex | *Ae. vexans* complex |
| Satara 2 | - | *C. pipiens* complex |
| Satara 3 | *Ae. vexans* complex; *An. gambiae s.l.; An. squamosus* | - |
| Satara 4 | - | - |
| Shingwedzi 1 | - | *C. univittatus* complex |
| Shingwedzi 2 | - | - |
| Shingwedzi 3 | - | *C. univittatus* complex*; An. gambiae* complex*; An. pretoriensis* |
| Shingwedzi 4 | *Ae. vexans* complex | - |
| Punda Maria 1 | - | *C. theileri* |
| Punda Maria 2 | - | *C. theileri* |
| Punda Maria 3 | - | - |
| Punda Maria 4 | *C. theileri* | - |

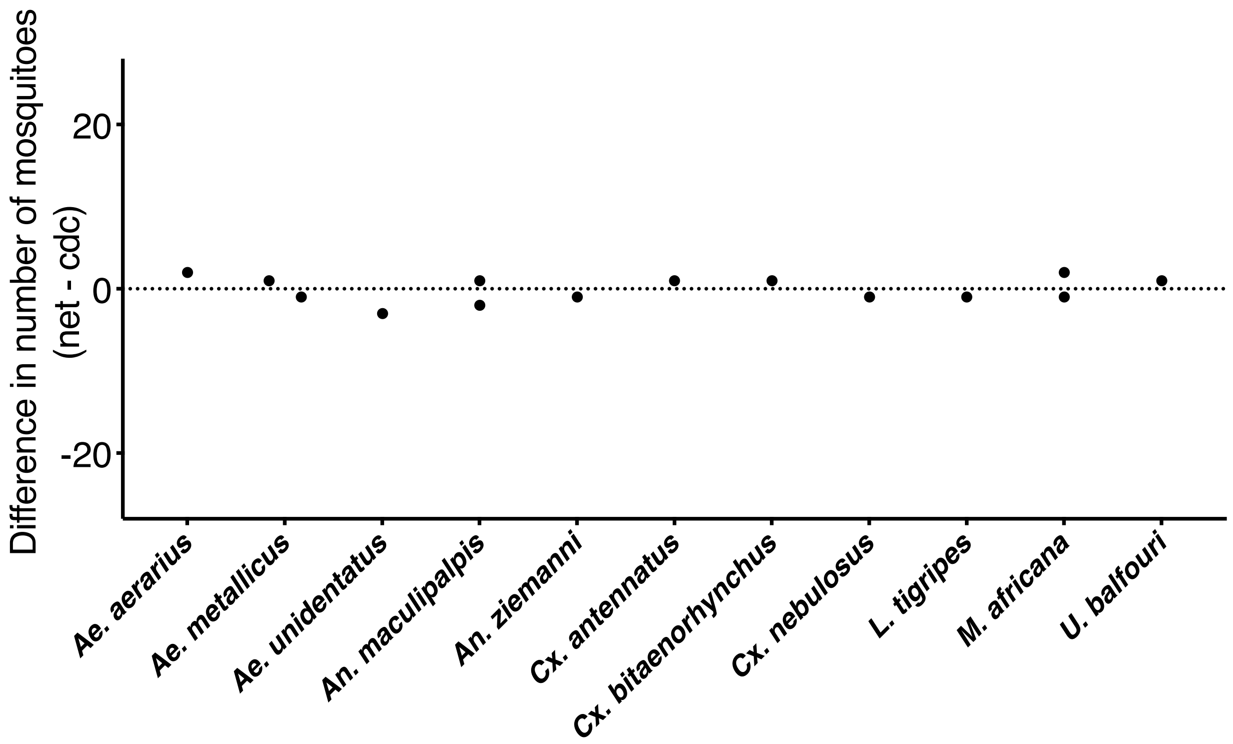

**Figure S3.** Species-specific trap preferences for the net *vs* CDC trap difference based on rare species not displayed in Fig. 3. Dots represent the difference in the number of mosquitoes collected in the net *vs* the CDC trap based on the total number of mosquitoes sampled across nights at each site.

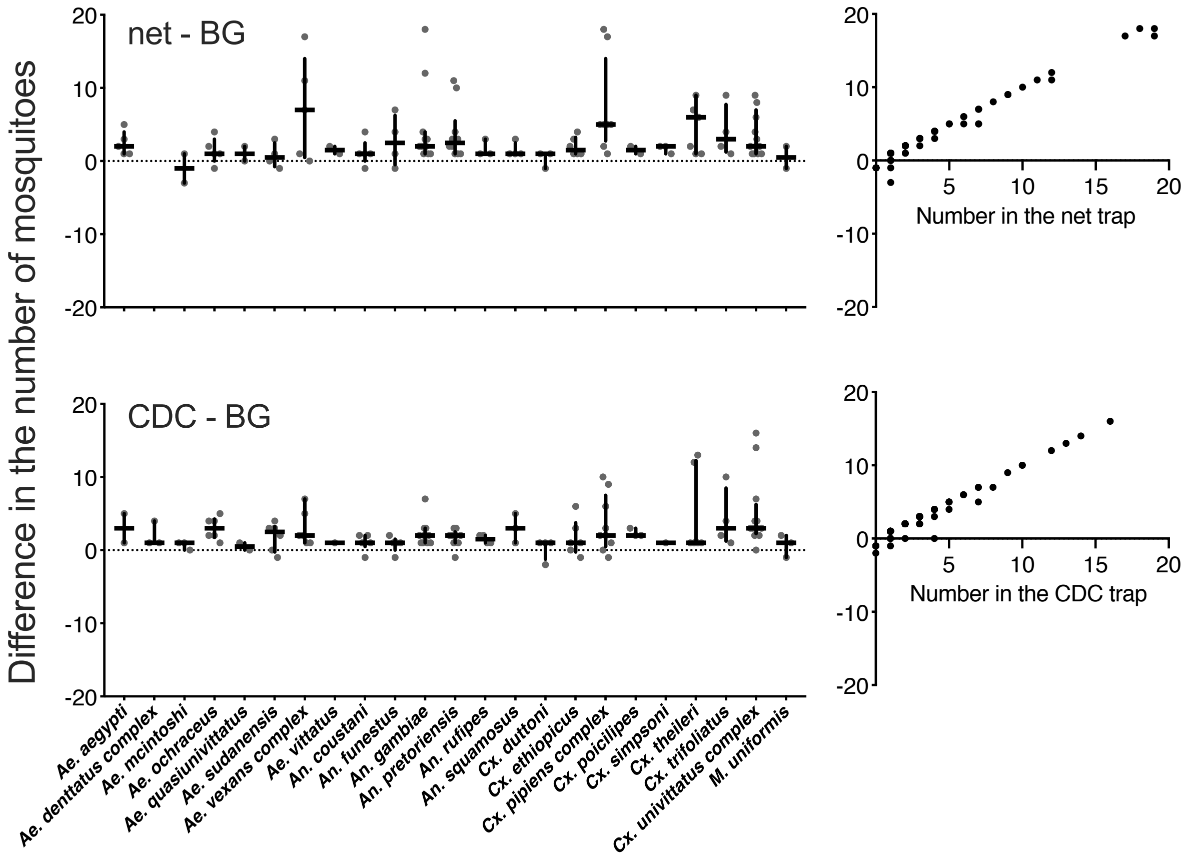

**Figure S4.** The net trap and the CDC trap caught higher numbers of mosquitoes (Fig. 3) and this pattern was not driven by any species or genus-specific trap bias (left figures) but by variation in the total number of the species collected (right figures).

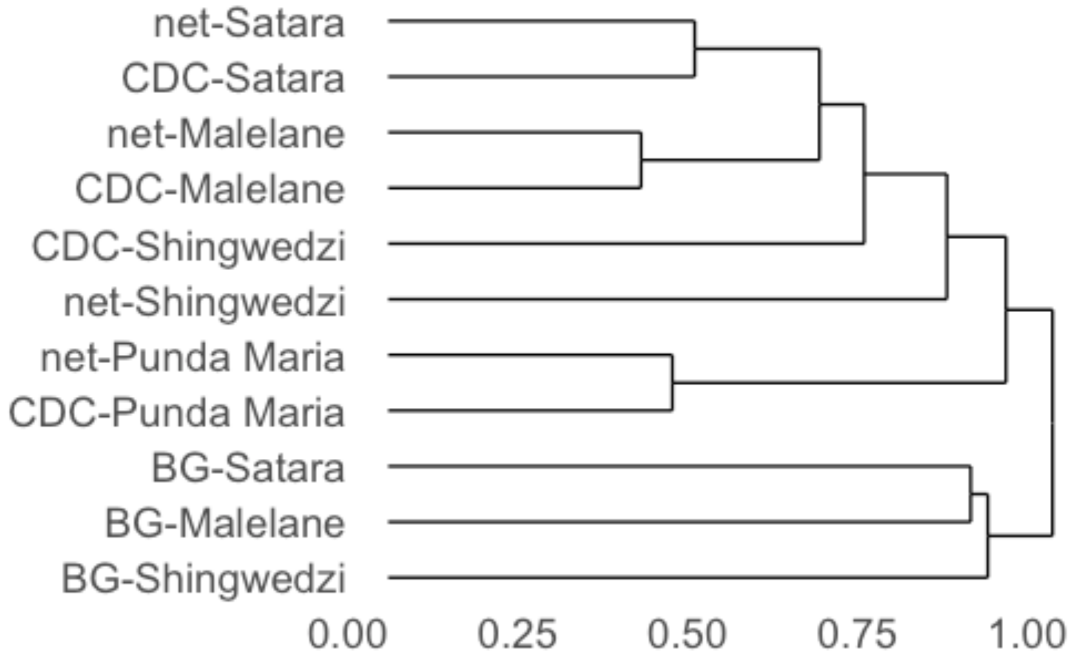

**Figure S5.** Dendrogram of species composition based on Bray-Curtis dissimilarity and the hierarchical clustering algorithm.
